# Increased activity in the somatosensory and insular cortex during the transition from acute to chronic neuropathic pain

**DOI:** 10.64898/2026.08.25.747057

**Authors:** Shayan Moosa, Erin D Murphy, Nikhil Gupta, W. Jeffrey Elias, Faraz Farzad, Chengsan Sun, Jaideep Kapur, Suchitra Joshi

## Abstract

Pathophysiological mechanisms underlying the transition from acute to chronic neuropathic pain remain incompletely understood. The somatosensory and insular cortices are key cortical components of the pain matrix. We examined changes in activation of these cortical regions during the transition from acute to chronic neuropathic pain.

The right sciatic nerve was ligated in activity reporter TRAP mice using standard procedures. Mechanical allodynia was confirmed after CCI or sham surgery using von Frey monofilaments applied to the hind paws. To label active neurons, 4-hydroxytamoxifen was administered to separate cohorts at 1, 3, and 6 weeks following nerve ligation. Passive tissue clearing of brain sections and confocal imaging was used to assess active neurons.

Progressive reduction of ipsilateral hind paw in CCI mice indicated mechanical allodynia development. CCI mice showed robust neuronal activation in the bilateral somatosensory and insular cortices. The somatosensory cortical activation peaked at 3 weeks post-CCI, whereas insular cortical activity increased during the transition from acute to chronic neuropathic pain.

These studies revealed that CCI induced progressive mechanical allodynia and distinct temporal patterns of cortical neuronal activation, with transient peak neuronal activity in the somatosensory cortex and sustained, increasing activation in the insular cortex during acute-to-chronic pain transformation.

## INTRODUCTION

Chronic pain remains a pervasive and debilitating condition, affecting millions worldwide and posing significant challenges for effective treatment. Approximately 20% of adults in the USA are estimated to suffer from chronic pain ^1,2^, with an accompanying annual cost of medical care ranging between $560 and $635 billion, according to the Institute of Medicine report Relieving Pain in America ^3^. Lack of sleep, anxiety, depression, and mood disturbances are some of the comorbidities of chronic pain that also lower the quality of life of the patients ^4^. Furthermore, the prescription of opioid drugs to alleviate chronic pain can lead to addiction ^5^.

Injuries or lesions to the peripheral nervous system are some of the common causes of neuropathic pain, besides neuropathies due to inflammatory disorders, diabetes, and cancer therapies ^6^. The development of allodynia and hyperalgesia indicates central sensitization, which is associated with molecular, synaptic, cellular, and network-level alterations in the pain matrix ^7^. Among these components, the somatosensory and insular cortices are especially important. The somatosensory cortices process the discriminative aspects of pain including location, intensity, and quality ^8,9^. In contrast, the insular cortex is important for the affective aspects of pain ^10–12^. How the engagement of these two cortical regions evolves during acute-to-chronic pain transformation remains unclear. In particular, the temporal dynamics of cortical neuronal activation during the transition from acute to chronic neuropathic pain have not been systematically characterized at cellular resolution. Understanding these network alterations is critical for revealing key regions of the pain matrix involved in maladaptive plasticity.

Chronic constriction injury (CCI) is a commonly used in vivo model of neuropathic pain to investigate the process of central sensitization ^13^. Using a CCI model of the sciatic nerve, we investigated changes in neuronal activation within the somatosensory and insular cortices during neuropathic pain chronification in a cross-sectional time-course study across independent cohorts. By combining behavioral assessment of hind paw mechanical allodynia with activity-dependent neuronal labeling in *T*argeted *R*ecombination in *A*ctive *P*opulations (TRAP) mice, we aimed to define spatial and temporal cortical activation patterns associated with the progression from acute to chronic neuropathic pain. Changes in neuronal activity maps will offer critical insights into how pain-related signals are processed, amplified, and sustained within the nervous system.

## MATERIALS and METHODS

### Animals

Adult (9- to 13-week-old) male and female TRAP mice were used in these studies ^14,15^. The studies were performed under a protocol approved by the Institutional Animal Care and Use Committee (IACUC). The animals were maintained on a 12-hour light-dark cycle (lights on at 6 AM and off at 6 PM) and had *ad libitum* access to food and water. Four to five mice of the same sex were housed together. Table 1 lists the sex of the mice used in the studies.

**Table 1:** Sex of the mice used in the study. M= males and F=females.

|  | Sham | CCI |
| --- | --- | --- |
| Acute | 1M 2F | 2M 2F |
| Subacute | 2M 1F | 2M 2F |
| Chronic | 1M 2F | 2M 1F |

### Unilateral chronic constriction injury (CCI)

The TRAP animals were randomized to sham surgery or CCI surgery and for the acute, subacute, and chronic time points. The sciatic nerve was constricted under isoflurane anesthesia to induce peripheral neuropathy. Briefly, an incision was made at the right mid-thigh of the anesthetized mouse, and the nerve was separated from the surrounding tissue. The constriction injury was performed proximal to the trifurcations with three loose ligatures at 1 mm intervals with a 5-0 silk thread to restrict but not completely halt blood flow to the area. The surrounding muscles were then rinsed with saline solution, and the incision was sutured closed. In addition, nine sham surgery mice were included in the study. These mice underwent surgery to expose the nerve, but no constriction was performed.

To differentiate the immediate, intermediate, and chronic effects of the nerve ligation, the mice were divided into three groups, and 4-hydroxytamoxifen (4OHT, 50 mg/kg in corn oil, subcutaneous) was administered at one, three, or six weeks after the sham or nerve ligation surgery. Since we wanted to evaluate changes in the basal neuronal activation, this injection was performed 24 hours after the paw withdrawal testing in the respective groups.

### Transcardial perfusion and passive tissue clearing

The animals were transcardially perfused six days after 4OHT administration, first with 10-15 ml of phosphate-buffered saline (PBS) followed by 40 ml of 4% paraformaldehyde containing 1% acrylamide ^16^. After overnight fixation in the same solution, the tissue was processed for passive tissue clearing ^16^. 200 *μ*m-thick horizontal brain slices were incubated overnight in 1% acrylamide containing 0.25% VA-044 at 4°C. The slices were then incubated in the same solution at 37°C for 2 hours, and the polymerizing solution was removed with 3 × 10-min PBS washes. The slices were then incubated overnight in PBS containing 8% SDS for 37°C tissue clearing. After 3 × 10-min PBS washes, the slices were counterstained with DAPI to label nuclei and mounted in a refractive index-matching solution.

### Imaging and quantification

The slices were imaged on a Nikon Eclipse Ti microscope under a 10x 0.75 NA lens. 10 *μ*m-thick optical sections at 1024 × 1024-pixel resolution were obtained with 550 nm excitation for tdTomato.

The images were analyzed using Imaris (Bitplane) software. Briefly, the region of interest (ROI) was outlined using the Paxinos mouse brain Atlas ^17^. The built-in “spots” function was used to count tdTomato-expressing cells. The number of active cells from the dorsal and ventral agranular insular areas (AID and AIV, respectively) was counted together. The tdTomato-positive neurons from the dysgranular and granular insular cortices (DI and GI, respectively) were counted together. Because the number of sections encompassing the ROI varied between animals, the data are expressed as the number of active neurons per hemisphere per slice.

### Evaluation of hind-paw mechanical thresholds using von Frey monofilaments

The testing was performed as described before ^18^. Briefly, the animals were habituated to Plexiglass boxes (5H × 5H × 7H) with wire-mesh floors. The testing was performed after 60 minutes of acclimatization. The mid-plantar surface of each hind paw was tested with von Frey filaments presented in ascending order, starting with the 0.4 g filament. The filaments were presented perpendicular to the paw until a slight buckling occurred and held for 3 seconds. There was a 10-second gap between each filament presentation. A quick withdrawal of the paw and/or its licking marked a positive response. Each filament was presented six times, and a filament that produced a positive response for 50% or more presentations was considered the threshold.

### Statistical analysis

The data were analyzed using GraphPad Prism software 10.4.1. The normality of the data was checked, and parametric or non-parametric tests were accordingly used to compare the data.

## RESULTS

### Development of mechanical allodynia

The TRAP mice were randomly divided into six cohorts, and their hind paw withdrawal thresholds were evaluated before and at 1, 3, or 6 weeks after the CCI or sham surgery to assess for changes in pain threshold (Fig. 1A). The left and right paw withdrawal thresholds in the sham surgery animals were similar and remained stable for 6 weeks of testing (Fig. 1B, Table 2). On the other hand, the withdrawal threshold in the CCI mice differed following ligation of the right sciatic nerve (Fig. 1B, Table 2). The pre-surgery thresholds in these animals were comparable; however, there was a sustained reduction in the right paw withdrawal threshold (ligated side) in CCI mice. The withdrawal threshold of the left paw (non-ligated side) was significantly higher at 6 weeks than at baseline (pre) and at 1-week post-surgery. The sustained lowering of the injured paw’s pain threshold demonstrated allodynia, as previously reported ^19^.

**Figure 1:**
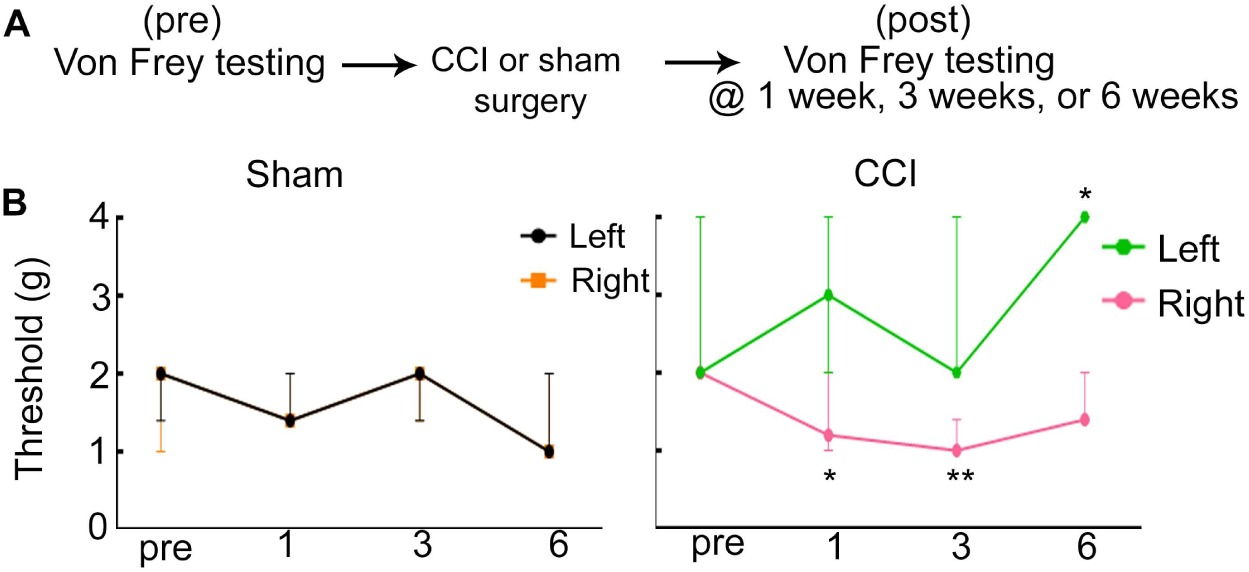
A reduction in the hind paw mechanical sensitivity indicative of the development of chronic neuropathic pain. **(A)** Experimental schematic. The left and right paw withdrawal thresholds were determined, and the mice were exposed to right sciatic nerve ligation (CCI) or a sham surgery in which the skin and muscles were opened and sutured back without injury to the sciatic nerve. **(B)** The paw withdrawal thresholds in the cohort of animals over the course of 6 weeks in the sham and CCI mice. Median and 95% CI are plotted. * p<0.05, ** p<0.005. Please refer to Table 2 for pairwise post-hoc Tukey’s multiple comparison tests.

**Table 2:**
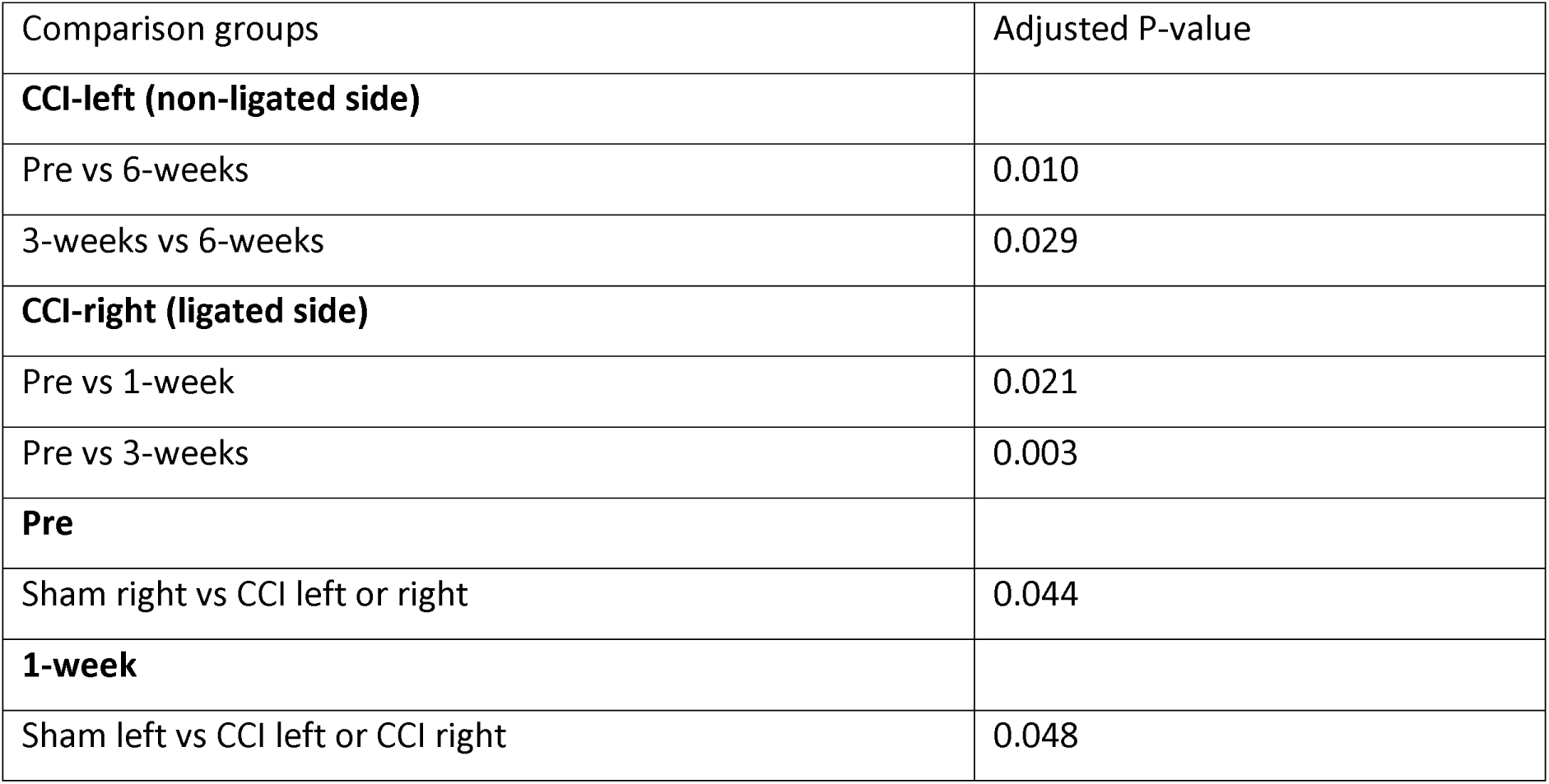
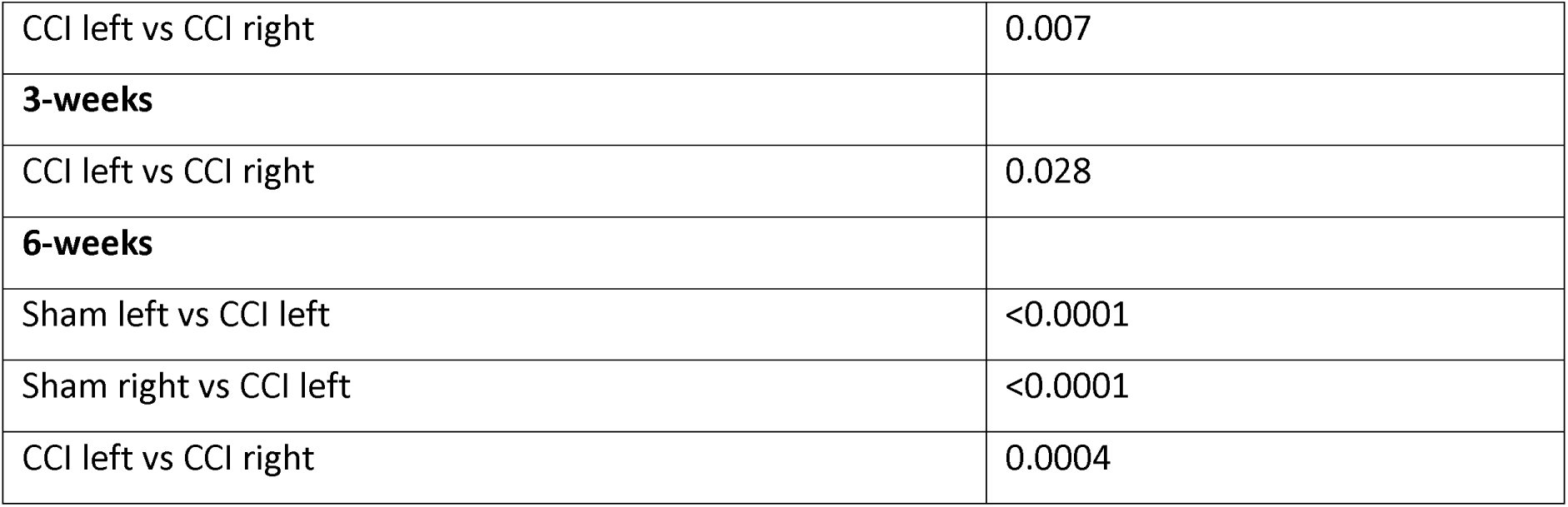
A summary of statistical comparisons of the pain threshold during the course of pain chronification in sham and CCI animals. Two-way ANOVA column factor (sham and CCI, left and right paw withdrawal threshold) p<0.0001, and interaction between the groups during the 6 weeks of the testing, p=0.0043. The post-hoc Tukey’s comparisons that yielded significant p values are listed below. None of the comparisons within the sham group were significant.

| Comparison groups | Adjusted P-value |
| --- | --- |
| **CCI-left (non-ligated side)** |  |
| Pre vs 6-weeks | 0.010 |
| 3-weeks vs 6-weeks | 0.029 |
| **CCI-right (ligated side)** |  |
| Pre vs 1-week | 0.021 |
| Pre vs 3-weeks | 0.003 |
| **Pre** |  |
| Sham right vs CCI left or right | 0.044 |
| **1-week** |  |
| Sham left vs CCI left or CCI right | 0.048 |
| CCI left vs CCI right | 0.007 |
| **3-weeks** |  |
| CCI left vs CCI right | 0.028 |
| **6-weeks** |  |
| Sham left vs CCI left | <0.0001 |
| Sham right vs CCI left | <0.0001 |
| CCI left vs CCI right | 0.0004 |

### Somatosensory cortical neuronal activation

The somatosensory cortex is the final node in the sensory and ascending pain pathways. To obtain insights into changes in neuronal activation during pain chronification, we evaluated its activation at 1, 3, and 6 weeks after ligation or sham surgery using activity-reporter TRAP mice, in which neurons active during the 60-90 minutes prior to 4OHT administration express the reporter protein tdTomato. There was prominent tdTomato expression in the somatosensory cortex of CCI animals, whereas the sham mice had sparse tdTomato expression (Fig. 2). The processes of active cells were also tdTomato-labeled (Fig. 2A, arrowheads), and their morphology was similar to that of the neurons. The labeling density appeared comparable between sham and CCI mice at one week. The CCI mice had many more labeled neurons at three and six weeks compared to sham animals. The labeling intensity appeared strongest at three weeks. The labeled neurons in CCI mice were concentrated in two bands across the cortex, corresponding to putative layers 4 and 6 at three weeks following nerve injury. In contrast, in the sham mice, the labeled neurons were distributed throughout the cortical thickness.

**Figure 2:**
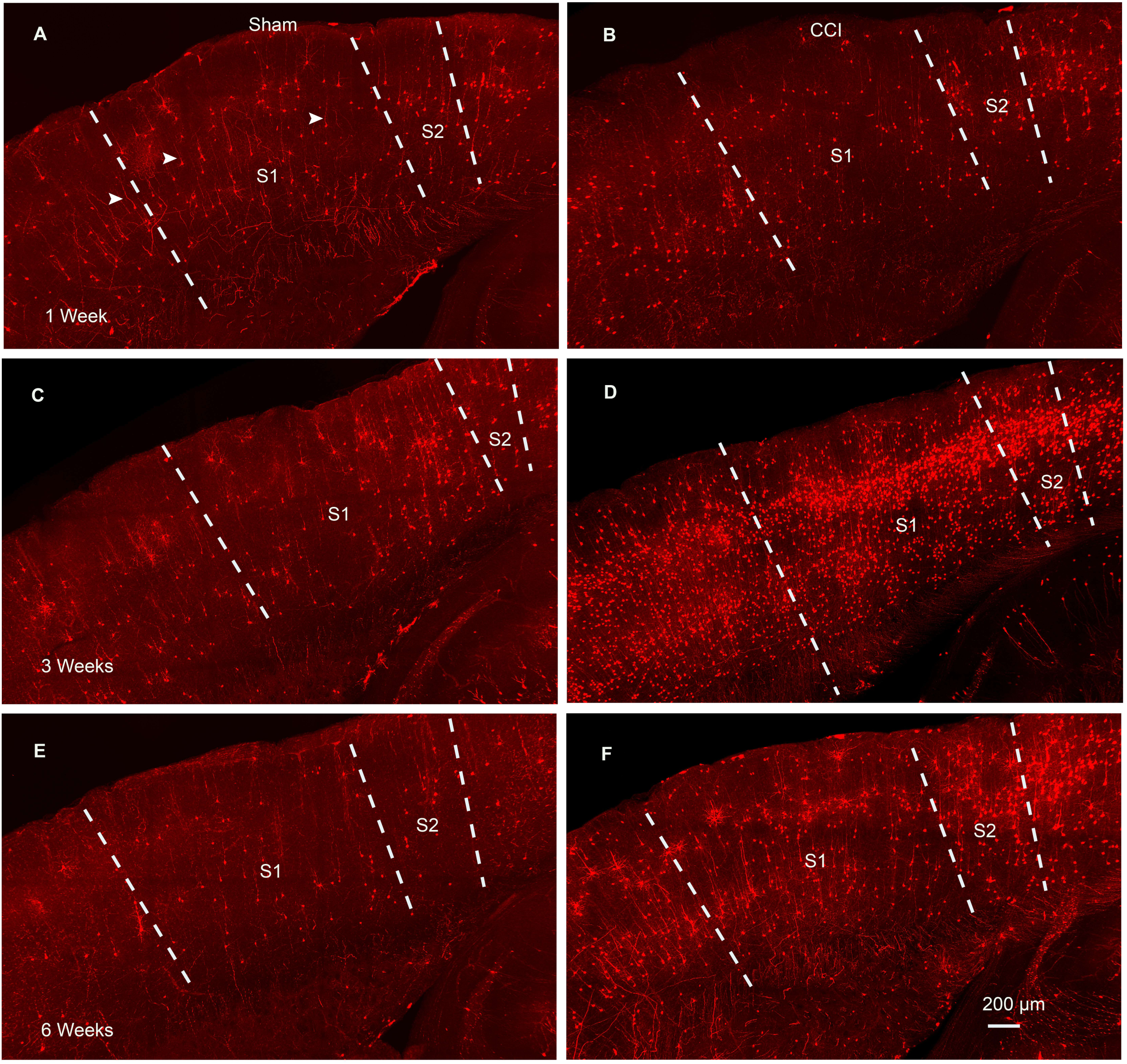
The activation of the somatosensory cortex during the development of chronic neuropathic pain. **(A, C, E)** tdTomato labeling in the somatosensory cortex of the sham surgery mice at one, three, and six weeks post-surgery, respectively. Arrowheads mark a few of the labeled processes. **(B, D, F)** tdTomato labeling in the somatosensory cortex of the CCI mice at one, three, and six weeks following injury, respectively. The dashed lines indicate the approximate boundaries of S1 and S2. Scale bar = 200 μm.

We counted the number of tdTomato-labeled cells. Within the CCI animals, the number of labeled cells in the primary and secondary somatosensory cortices (S1 and S2, respectively) was highest at three weeks (Fig. 3A, 3D). In the sham group animals, more labeled cells were present at one week, likely an acute effect of the surgery, than at three and six weeks (Fig. 3B, 3E). The group comparisons revealed that the neuronal activation in the CCI and sham animals was comparable at one-week post-surgery (Fig. 3C, 3F). The number of active neurons in the CCI mice was 6-10 times more than that in sham mice at three weeks and 5-7 times more at six weeks (Fig. 3C, 3F). Thus, allodynia development in CCI animals was associated with activation of primary and secondary somatosensory cortical neurons, which was first observed 3 weeks after surgery.

**Figure 3:**
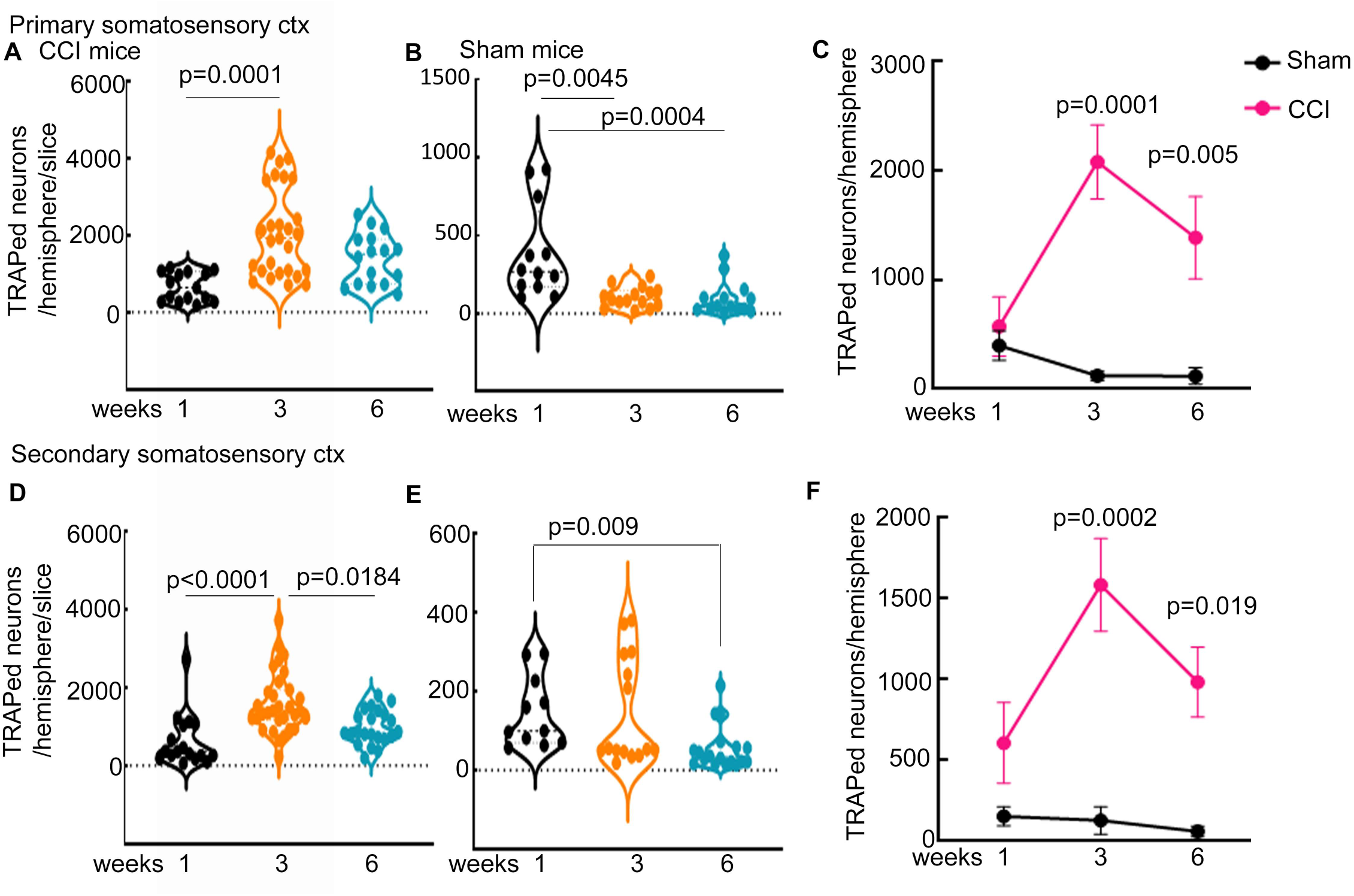
Somatosensory cortical activation during chronic neuropathic pain. **(A and B)** The number of tdTomato-positive (TRAPed) cells in the primary and secondary somatosensory cortices from individual slices from CCI and sham group animals at one (n=6 slices for CCI and n= 12 slices for sham), three (n=19 slices for CCI and n=14 slices for sham), and six (n=9 slices for CCI and n=15 slices for sham) weeks after the surgery. The number of animals from which the slices came is listed in Table 1. The data did not pass the normality test for the acute and chronic time points; hence the Kruskal-Wallis test was used for comparing all six groups, p=0.0003. The significant differences in the post-hoc pair-wise Dunn’s multiple comparison test are marked on the graph. **(C)** Mean and SEM of TRAPed primary somatosensory cortical neurons per hemisphere in sham and CCI animals at one, three, and six weeks after surgery. The number of replicates is the same as that listed in Table 1. Two-way ANOVA revealed significant differences between the sham and CCI animals, p=0.0003, F (1, 12)= 26.23. Pairwise post-hoc Tukey’s multiple comparison tests revealed significant differences between the sham and CCI animals at three and six weeks after surgery. **(D and E)** The number of TRAPed cells in the secondary somatosensory cortex in individual slices of CCI and sham animals, respectively (CCI: n=17 slices one week, n=31 slices three weeks, and n=22 slices six weeks and sham: n=11 slices for one week, n=15 slices three weeks, and n=18 slices six weeks). Comparison of the TRAPed cells from all six groups using the Kruskal-Wallis test revealed p<0.0001. The p-values, wherever significant, of pairwise post-hoc Dunn’s multiple comparisons are marked on the graph. **(F)** Mean and SEM of TRAPed secondary somatosensory cortical neurons at one, three, and six weeks after the sham or CCI surgery. The number of replicates is the same as that listed in Table 1. Two-way ANOVA revealed significant differences between the sham and CCI animals, p<0.0001, F (1, 14)= 29.42. The p-values of pair-wise post-hoc Tukey’s multiple comparison tests, wherever significant, are marked on the graph.

### Activation of the insular cortex

The insula plays a critical role in pain processing and serves as a central hub connecting ascending and descending pain pathways ^20^; its stimulation elicits pain ^21,22^, and its lesions alleviate pain ^23^. Hence, we also evaluated its activation during acute-to-chronic neuropathic pain transformation. We matched the horizontal brain sections with the Paxinos Atlas, and the areas were marked based on landmarks and the lack of layer IV in the agranular insular cortex.

The agranular insular cortex (AID/AIV) forms the rostral/anterior insular cortex, particularly critical for regulating emotional aspects of pain through circuitry also involving the anterior cingulate cortex and amygdala ^20^. We evaluated tdTomato expression in the AID/AIV in sham and CCI mice. The labeled cells were sparse in the sham mice but abundant in the CCI mice (Fig. 4). As seen in the somatosensory cortex, the processes of these cells were also labeled with tdTomato fluorescence, indicating their neuronal nature. In the AID/AIV of the CCI mice, there was a progressive increase in the number of labeled neurons from one to three to six weeks (Fig. 5A). In contrast, the sham mice had more active AID/AIV neurons at one-week post-surgery than at three and six weeks (Fig. 5B). Thus, the number of active AID/AIV neurons increased continuously in the CCI mice and decreased in the sham mice.

**Figure 4:**
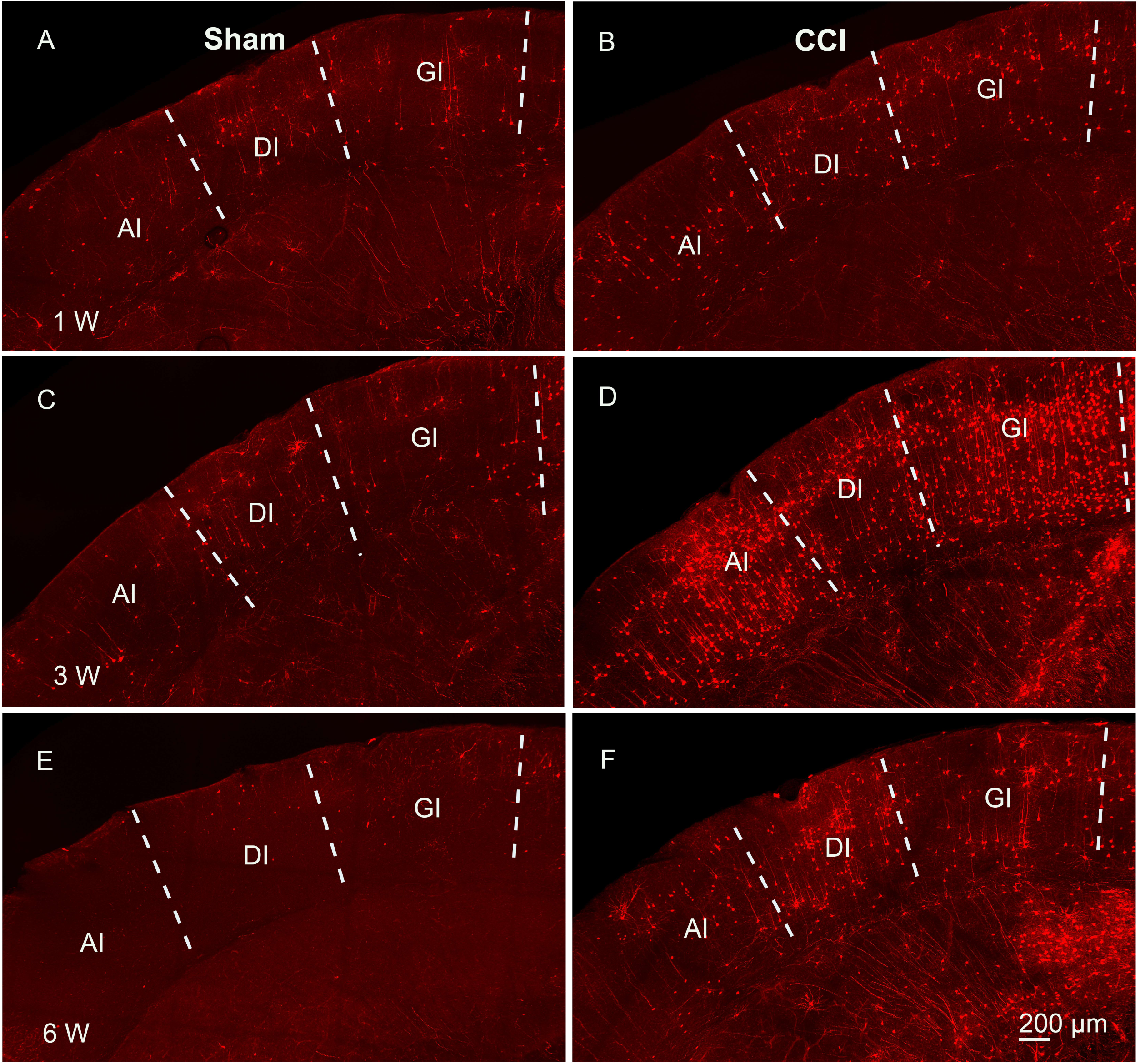
The activation of the insular cortex during the development of chronic neuropathic pain. **(A, C, E)** tdTomato labeling in the insular cortex of the sham surgery mice at one, three, and six weeks post-surgery, respectively. **(B, D, F)** tdTomato labeling in the insular cortex of the CCI mice at one, three, and six weeks, respectively, after the surgery. The dashed lines indicate the approximate boundaries of the agranular (AID/AIV), dysgranular (DI), and granular (GI) subdivisions of the insular cortex. Scale bar = 200 μm.

**Figure 5:**
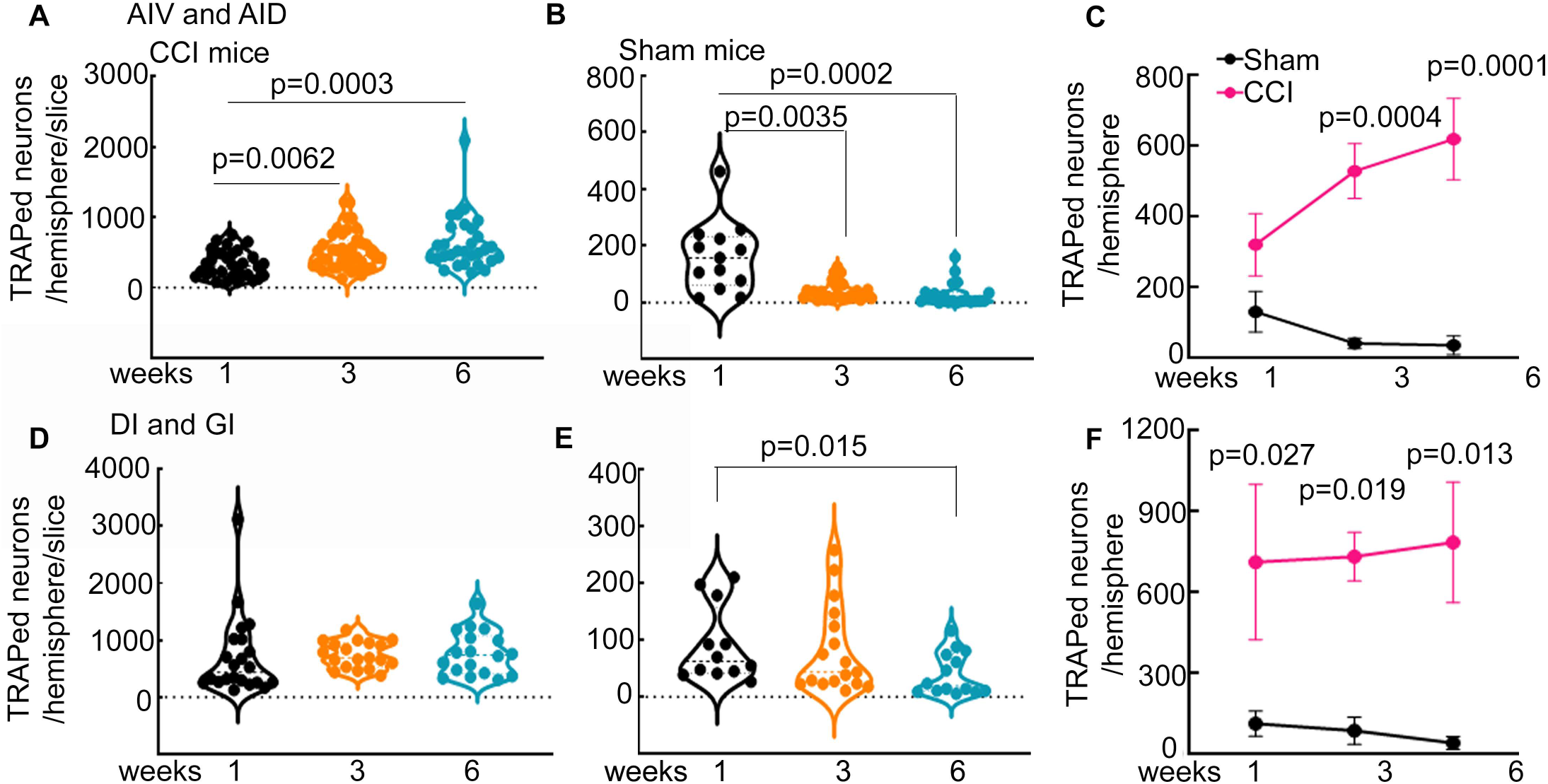
Activation of the insular cortex during neuropathic pain chronification. **(A and B)** The number of tdTomato-positive (TRAPed) cells in the agranular insular cortex (AID and AIV) per hemisphere from individual slices from CCI and sham group animals, respectively, at one (n= 32 for CCI and n=13 slices for sham), three (n=40 for CCI and n=20 for sham), and six (n=30 for CCI and n=20 slices for sham) weeks. **(C)** Mean and SEM TRAPed neurons per hemisphere in each animal in the two treatment groups at one, three, and six-week time points. The number of replicates is the same as that listed in Table 1. Two-way ANOVA revealed significant differences between the sham and CCI animals, p<0.0001, F (1, 14)= 45.6. The p-values of pair-wise post-hoc Tukey’s multiple comparisons, wherever significant, are marked on the graph. **(D, E)** The number of TRAPed neurons in the dysgranular and granular insular cortex in the CCI and sham animals, respectively (one week: n=22 CCI and n=12 sham; three weeks: n=19 CCI and n=11 sham; six weeks: n=18 CCI and n=14 sham). **(F)** Mean and SEM TRAPed neurons in the animals belonging to the two treatment groups at one, three, and six weeks. Two-way ANOVA p=0.0004, F (1, 14)= 21.2. The p-values of pair-wise post-hoc Tukey’s multiple comparisons, wherever significant, are marked on the graph.

The dysgranular and granular insular cortex (DI/GI) also had tdTomato-labeled neurons, sparser in the sham mice than in the CCI mice (Fig. 4). Within the CCI mice, the number of labeled neurons remained constant from one week to six weeks (Fig. 5D). In the sham groups, more neurons were labeled at one week than at six weeks (Fig. 5E). Finally, comparison of active neurons in sham and CCI mice revealed a significantly greater number of insular cortical neurons in CCI mice (Fig. 5C, 5F). The CCI mice had 2-5 times more active neurons than the sham mice at one, three, and six weeks (Fig. 5F). The data from granular, dysgranular, and agranular insular cortex taken together showed a progressive increase in active neurons during the development of chronic neuropathic pain.

### Laterality of neuronal activation

Although the injury was unilateral, TRAPed neurons were present in both hemispheres. We did not mark the ipsilateral hemisphere and were unable to distinguish between the two sides. To assess if both hemispheres had similar neuronal activation, we took the ratio of active neurons in one hemisphere to that in the other; a similar neuronal activation in both hemispheres would yield these ratios close to 1. We reasoned that if both hemispheres had a similar number of neurons, the ratio of active neurons in one to the other hemisphere would be close to 1. The distribution of active neurons across the hemispheres was similar in the sham animals across all the cortical regions evaluated, and it remained unaltered during the six weeks of evaluation (Fig. 6). The active neurons were also spread across both hemispheres of the primary and secondary somatosensory cortices of the CCI mice (Fig. 6A, 6B). On the other hand, the distribution of active neurons across the two hemispheres was uneven in the insular cortex. One hemisphere consistently had more active neurons than the other hemisphere (Fig. 6C, 6D).

**Figure 6:**
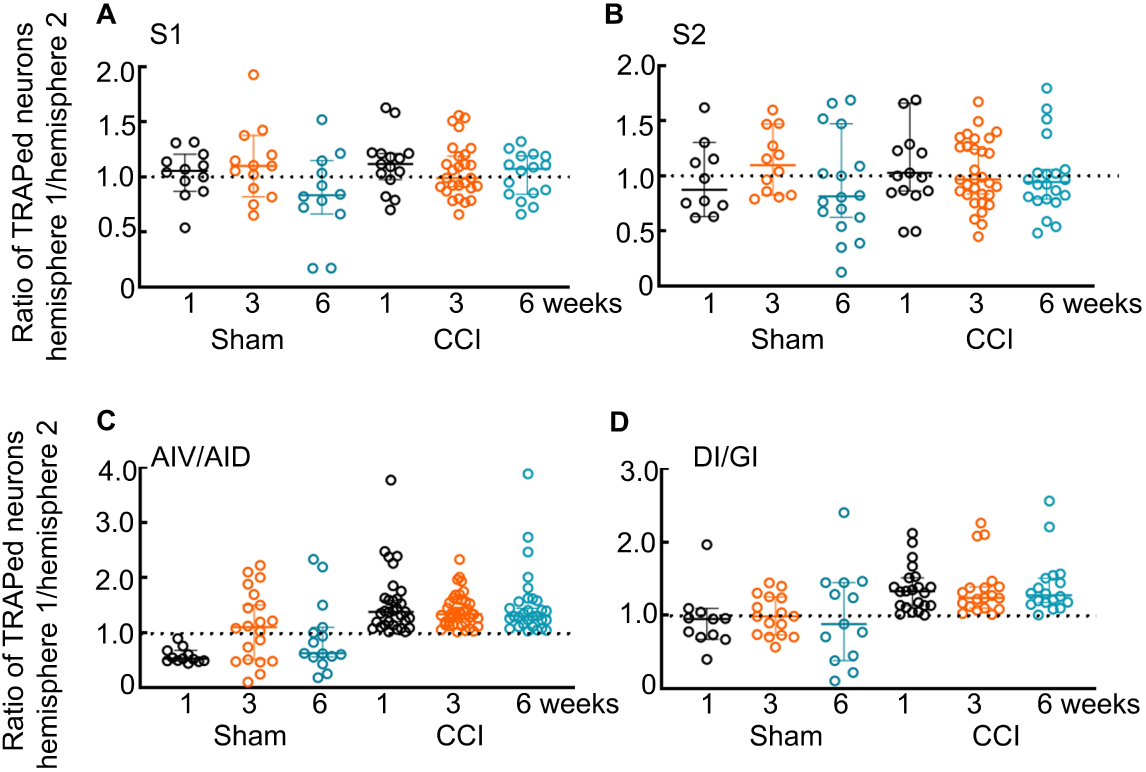
Laterality of neuronal activation. **(A, B)** The ratio of active somatosensory cortical neurons in one hemisphere to that in the other hemisphere. The ratios were determined for each slice analyzed in the sham and CCI mice at one, three, and six weeks. The dotted line marks the ratio of 1 indicative of equal distribution of active neurons in the two hemispheres. **(C, D)** The ratio of active neurons in the two hemispheres in the insular cortex. While the ratios in the sham animals were distributed evenly on either side of 1, those in the CCI animals were always above 1, suggesting that the number of active neurons in one hemisphere was always higher than that in the other. The number of slices is the same as in Figures 2 and 5.

## DISCUSSION

The chronic constriction injury mouse model predictably caused a sustained lowering of the ipsilateral paw withdrawal threshold (indicative of mechanical allodynia), and this behavioral change correlated with the bilateral activation of the primary and secondary somatosensory and insular cortices. The activation of the somatosensory cortex peaked at three weeks after the CCI and then decreased while the insular cortical activity continued to increase. This suggests that central sensitization occurs first in the somatosensory cortex and later in the anterior insula.

The paw withdrawal threshold lowered on the ipsilateral side of the sciatic nerve injury. These findings agree with prior studies that have also reported hypersensitivity to mechanical and thermal stimuli in this model ^24,25^.

c-Fos, an immediate early gene, is widely used as a proxy for neuronal activation, and we used c-Fos to drive tdTomato reporter expression in the activated neurons. We found tdTomato expression in the somatosensory (S1 and S2) and insular cortices of CCI mice; the tdTomato labeling was stronger in the CCI mice compared to the sham-operated mice. Thus, there was a sustained engagement of central pain-processing circuits during chronic neuropathic pain development. Increased activation of the insular cortex was evident a week after CCI surgery, and it remained high throughout the study duration. The activation of the somatosensory cortices emerged later. A growing body of evidence indicates that neuropathic pain is maintained by widespread cortical plasticity rather than peripheral or spinal mechanisms. The sustained cortical activation seen here supports this notion.

Interestingly, the somatosensory cortical activation was bilateral, and the density of activated neurons was comparable between the hemispheres. Insular cortical activation on the other hand, appeared to be stronger in one hemisphere than the other. Nerve injury-induced central sensitization can lead to recruitment of homologous cortical regions in both hemispheres through interhemispheric connectivity and thalamocortical network reorganization. Bilateral activation of the anterior cingulate cortex following nerve injury is also seen in another study ^26^. In contrast, other studies have found ipsilateral activation of the spinal dorsal horn ^24,25^, and that of parabrachial and locus coeruleus regions of the brainstem ^25,27^. We did not distinguish neuronal from non-neuronal tdTomato expression. However, the labeling of processes and projections of the activated cells suggests that a majority of them were neurons.

Stimulation of secondary somatosensory and insular cortices in humans triggers pain ^28^. Electrical stimulation of the insula produces pain sensation in humans ^28,29^, and lesions within the insula have been shown to alter pain perception ^12,30–32^. The insular cortex is also a key hub in the affective and interoceptive aspects of pain ^12^. Converging evidence from both animal models and human studies indicates that the insula exhibits increased activity and synaptic potentiation in neuropathic conditions. For instance, functional imaging in rodent nerve injury models has revealed bilateral activation of the insular and somatosensory cortices during neuropathic pain states. Moreover, molecular and electrophysiological studies demonstrate enhanced excitatory transmission and plasticity within the insula following peripheral nerve injury, supporting its role in central sensitization. The involvement of the insular cortex is further supported by lesion and intervention studies. Disruption of insular subregions, particularly the caudal granular insular cortex, can attenuate allodynia associated with neuropathic pain ^33^, indicating that this region is not merely responsive but causally involved in pain maintenance. Thus, increased activation of the insula likely reflects both heightened nociceptive processing and engagement of circuits that sustain chronic pain. While the activation of the somatosensory cortices was bilateral, the ratio of active insular cortical neurons in one hemisphere to those in the other consistently stayed above one, suggestive of a degree of laterality. The ipsilateral paw withdrawal thresholds were lowered, and hence we anticipate the contralateral insular cortex to contain more active neurons. However, in this study, we did not distinguish the ipsilateral and contralateral hemispheres, and follow-up studies are needed to evaluate if this is the case.

Direct evidence that the insula is a potential therapeutic target is being investigated by our group. In a pilot study of six patients with epilepsy undergoing intracranial monitoring for seizure localization, direct electrical stimulation of the anterior insula significantly raised the heat pain threshold, reduced laser-evoked pain intensity, attenuated nociceptive-specific laser-evoked potentials, and altered ongoing intracranial EEG band power. ^34^ We are now studying neuromodulation of the anterior insula in patients with refractory neuropathic pain. (NCT05404581) Our observation that insular activation rose progressively and remained elevated while somatosensory cortical activation peaked and then declined suggests that the insula may be a key region involved during the transition from acute to chronic pain.

Some limitations should be considered. The c-Fos-driven tdTomato expression provides only a snapshot of neuronal activation, and it cannot resolve circuit-level dynamics or causal relationships. Also, the current study did not distinguish between the activation of excitatory and inhibitory neuronal populations. Future studies using cell-type-specific approaches, including calcium imaging and chemogenetic or optogenetic manipulation of neuronal ensembles, would help clarify the precise contributions of these cortical regions. Such manipulations of neuronal activity are known to regulate pain thresholds. For example, optogenetic inhibition of the anterior cingulate cortex improves mechanical and thermal pain thresholds and offers pain relief in nerve injury-induced neuropathic pain^35^. Chemogenetic silencing of progesterone receptor-expressing somatosensory cortical neurons alleviates migraine pain ^36^.

The current study focused on changes in the somatosensory and insular cortical activation in neuropathic pain chronification. Other regions of the pain matrix are also likely to be activated, and future studies evaluating the activation of sensory thalamus and brainstem nuclei will be critical. Alterations in excitability, synaptic plasticity, and network reorganization have been noted in other cortical areas in chronic neuropathic pain. For example, one to two weeks after the CCI, the excitatory drive on interneurons in L5 of the anterior cingulate cortex is reduced simultaneously with increased excitability of the principal neurons of this layer and reduced inhibitory drive on these neurons ^37^. Nerve injury also increases spine density and complexity in L2/3 of the medial prefrontal cortex ^38^. M1 receptor internalization leading to attenuation of the cholinergic excitatory drive on L5 neurons of the medial prefrontal cortex is seen in neuropathic pain ^39^. Evaluation of synaptic plasticity of the activated neurons would also provide additional insights into molecular mechanisms underlying the transformation of a normal network to a hyperexcitable circuit.

In conclusion, increased bilateral activation of the somatosensory and insular cortices following CCI supports a model in which neuropathic pain is maintained by distributed cortical hyperactivity and plasticity, encompassing both sensory and affective domains. These findings further highlight the insular cortex as a potential therapeutic target and underscore the importance of considering brain-wide network changes in chronic pain.

## Data availability

The data are available from the corresponding author.

## Acknowledgements

We thank John Williamson for critical comments. SJ is supported by R21 NS 140790-01A1.

## Author contributions

Data collection: SM, FF; data analysis: SM, EM, NG, SJ; Study conceptualization and manuscript preparation: all authors

